# Comparative analysis of temporal transcriptome reveals the relationship between pectin degradation and pathogenicity of defoliating *Verticillium dahliae* to Upland cotton (*Gossypium hirsutum*)

**DOI:** 10.1101/2020.10.02.323402

**Authors:** Fan Zhang, Jiayi Zhang, Wanqing Chen, Xinran Liu, Cheng Li, Yuefen Cao, Tianlun Zhao, Donglin Lu, Yixuan Hui, Yi Zhang, Jinhong Chen, Jingze Zhang, Alan E. Pepper, John Z. Yu, Shuijin Zhu

**Author notes:** Department of Biology, College of Science, Texas A&M University, College Station, Texas, United States of America (Fan Zhang, Alan E. Pepper). Corresponding author: (A.E.P.), (J.Z.Y.), and (S.J.Z).

## Abstract

Verticillium wilt (VW), caused by *Verticillium dahliae* Kleb., is a major plant disease that causes heavy annual losses around the world, especially in Upland cotton (*Gossypium hirsutum*). The disease-causing pathogen can be classified into defoliating (D) and non-defoliating (ND) pathotypes based on the induced symptoms. At present, little is known about the complex mechanisms of fungal pathogenicity and cotton resistance to it. Comparative analysis of temporal transcriptome was performed on two *V. dahliae* strains, *Vd_086* (D) and *Vd_BP_2_* (ND), at key development stages (hyphal growth, microsclerotia production, and spore germination) to reveal the functional process on plant defoliation and death. Differentially expressed gene (DEG) analysis revealed a strong correlation between cell wall protein kinase activities and the early pathogenicity of defoliating *Vd_086*. With weighted gene co-expression network analysis (WGCNA), six specific gene modules were correlated with the biological traits of the fungal samples. Functional enrichment with Gene Ontology (GO) and Kyoto Encyclopedia of Genes and Genomes (KEGG) pathways together with DEG analysis revealed six pectin degrading enzymes including *Polygalacturonase gene 1 (PG1)*, *Pectate lyase gene (PEL)* and *Pectinesterase gene 1 (PME1)* expressed in the early development of *Vd_086* that may be related to the robust pathogenicity of this strain during the early invasion. The expression of four of these genes was verified by real-time quantitative reverse transcription PCR (qRT-PCR). In addition, we identified Mitogen-Activated Protein Kinase (MAPK) signaling “hub” genes that may regulate these pectinases. In a word, enhanced expression of pectin degradation enzymes is associated with the stronger pathogenicity of *Vd_086* than *Vd_BP_2_*, especially at early infection stages. The disease-causing capability is likely regulated by MAPK signaling genes. This study provides new insight into molecular mechanisms of the plant-pathogen interaction on the VW disease, facilitating more effective control measures against this pathogen, including molecular breeding for the VW-resistant cotton cultivars.

**Author summary:** Verticillium wilt (VW), caused by fungal pathogen *Verticillium dahliae* (*Vd*), is arguably the most devastating disease in cotton production for decades. Molecular biologists and plant breeders have been working hard to identify host plant resistant genes for many years but have met with little success due to the large complex genome of cotton. The *V. dahliae* strains are grouped in two pathotypes, of which defoliating (D) strains cause total leaf loss of infected cotton plants and non-defoliating (ND) strains do not. Comparative transcriptome analysis of D strain *Vd_086* and ND strain *Vd_BP_2_* identified the candidate genes and molecular mechanisms related to the *Vd* pathogenicity. Besides the difference in pathogenicity, these strains are distinguishable by the rate of hyphal elongation, microsclerotia production, and spore germination. With these phenotypes, transcriptome sequencing of both strains was performed at the three growth phases. By the combination of comparative transcriptomic differentially expressed gene (DEG) analysis and weighted gene correlation network analysis (WGCNA), cell wall-associated pectinase genes were found to be active at hyphal elongation stage of the *V. dahliae* pathogen and ribosome-related processes were activated for microsclerotia production. Gene modification processes were activated with many protein kinases at spore germination stage that for the next infection cycle. Furthermore, four pectinases in the pentose and glucuronate interconversion (PGI) pathway were identified and verified as highly expressed in the D strain with strong pathogenicity to Upland cotton (*Gossypium hirsutum*). Our results provided evidence in support of the hypothesis that stronger early pathogenicity of the D strain is resulted from greater plant cell wall pectin degradability. Transcription factors (TFs) and “hub” module genes were identified in searching of protein interaction for possible regulators of the recognized pectinases. TFs involved in mitogen-activated protein kinase (MAPK) signaling pathway were shown to regulate not only hyphal processes but also the entire growth period of *V. dahliae*. This is the first study known to use module extraction techniques of WGCNA to identify differentially co-expressed genes between two fungal pathotypes of *V. dahliae* strains. The study provides new insights into molecular mechanisms of the plant-pathogen interaction and may lead to molecular breeding for resistant cotton cultivars to effectively control this devastating disease.

## Introduction

Verticillium wilt (VW) diseases in crop plants are spreading worldwide, especially where susceptible crops are grown. They affect agricultural economies of most countries to various extents ^[1]^. Major crops affected by VW diseases include cotton, potato, tomato, strawberry, eggplant, olive, sunflower, tobacco, and many herbaceous and woody perennials ^[2, 3]^. Among all these plant species, cotton (*Gossypium* spp.), the leading natural fiber crop, suffers the most severe economic losses ^[4]^. Of four cultivated cotton species, the disease primarily affects Upland cotton (*Gossypium hirsutum* L) which produces approximately 95% world’s natural fiber.

The causal agent is *Verticillium dahliae* Kleb, a soil-borne fungus capable of infecting plant roots and colonizing the vascular tissues throughout growth season, which was first reported more than a century ago ^[5]^. In general, *V. dahliae* is classified into two pathotypes: defoliating (D) and non-defoliating (ND), based on induced symptoms ^[6]^. The D pathotype is highly virulent and usually causes a severe wilting syndrome including chlorosis, fall of green leaves (defoliation), vascular discoloration, and death of the plant. The ND pathotype does not cause such severe phenotype over the same infection period (Fig. 1A, B). Infected plants exhibit symptoms of decreasing in photosynthesis and increasing in respiration, resulting in a significant reduction of the plant biomass and heavy loss of yield ^[7]^. In addition to the different pathogenetic capabilities, the two strains show different growth phenotypes outside of the host, when cultured on Potato Dextrose Agar (PDA) plates (Fig. 1C). This pathogen has a broad host range of more than 400 plant species, and can survive extremely long periods of time in the soil as microsclerotia, a heavily melanized resting structure ^[8]^.

**Fig. 1.**
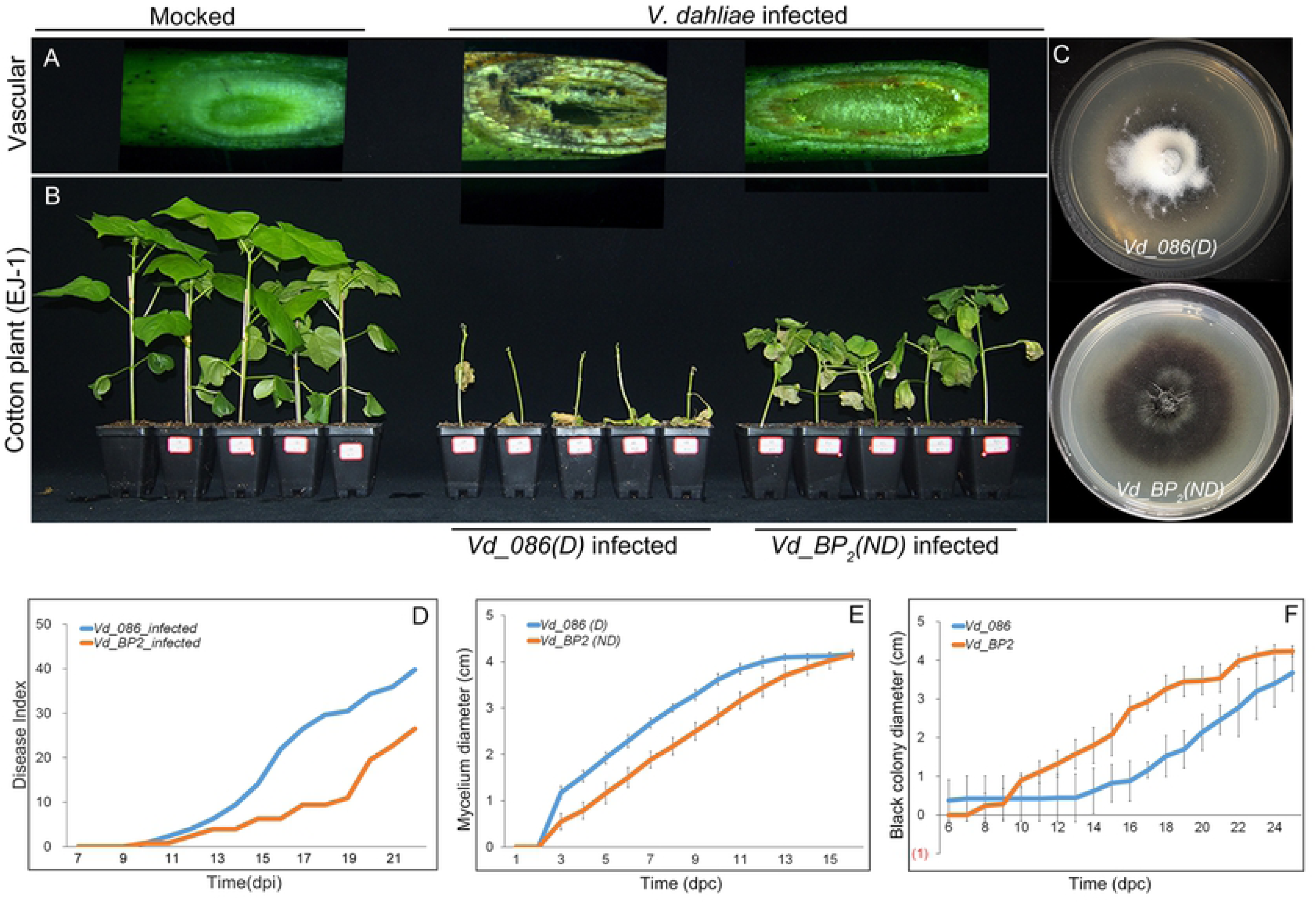
Disease symptoms of infected cotton plants and assessment of pathogen morphology. (A) Xylem vessel browning of cotton seedlings inoculated with distilled water, *Vd_086* (defoliating strain) spores, and *Vd_BP_2_* (non-defoliating strain) spores on the 21^st^ days post inoculation (dpi). (B) Disease symptoms of susceptible Upland cotton species EJ-1 with inoculation of distilled water, *Vd_086* spores, and *Vd_BP_2_* spores on the 21^st^ dpi. (C) Morphology of *Vd_086* and *Vd_BP_2_* strains with 20 days post incubation. (D) Disease progress curves of cotton plants inoculated by *Vd_086* and *Vd_BP_2_* spores with dpi. Disease index (DI) at each time point is calculated from 32 replicates with 5-grade disease index statistical method (see details in Methods). (E) Hyphal elongating progress by measurement of mycelium diameter of *Vd_086* and *Vd_BP_2_* with days post culturing (dpc) on PDA medium. Error bars represent standard deviations calculated from 6 - 10 replicates. (F) Black microsclerotia growing progress by measurement of black colony diameter of *Vd_086* and *Vd_BP_2_* with dpc on PDA medium. Error bars represent standard deviations calculated from 6 - 10 replicates.

Conventional management for cotton VW disease is an integrated approach that includes planting resistant cultivars, extending cultural practices ^[3]^, managing water and fertilizer ^[9]^, and biocontrol with fungicidal bacteria ^[10]^. Among these, breeding resistant varieties is considered the most effective and economical approach to controlling this disease. Currently, however, the breeding for the resistant cultivars has met with little progress due to the lack of highly resistant cotton germplasm ^[11]^. Hence, the research on molecular mechanisms of cotton VW disease has become a priority in recent years. Some plant resistance genes have been elucidated to play important roles in the prevention of VW diseases. The *non-expresser of pathogenesis-related genes-1* (*NPR1*), encodes a key regulator in the salicylic acid-mediated induction of plant systemic acquired resistance (SAR) ^[12]^. However, Parkhi et. al. (2010) demonstrated that the expression of *Arabidopsis NPR1* in transgenic cotton confers resistance to the ND isolates of *Verticillium dahliae* but not to the defoliating ones ^[13]^. As *G. hirsutum* is currently the most widely cultivated but is susceptible cotton, it is the focus of many studies on prospective VW resistant genes. Gao et. al. (2011) discovered that silencing *GhNDR1* (*G. hirsutum non-race-specific disease resistance-1*) and *GhMKK2* (*G. hirsutum MAP kinase kinase-2)* compromises cotton resistance to the infection by *V. dahliae* ^[14]^. The *G. hirsutum* TIR-NBS-LRR gene *GhDSC1* also mediates the resistance response associated with reactive oxygen species (ROS) accumulation by increasing expression of JA-signaling-related genes ^[15]^. Gong et. al. (2017) demonstrated that the *ribosomal protein L18 gene* (*GaRPL18*) plays a crucial role in resistance to cotton VW disease by regulating an SA-related signaling pathway. Yang et. al. (2018) revealed the receptor-like protein *Gbvdr6* confers plant resistance through regulation of the JA/ET and SA signaling pathways ^[16, 17]^. Recent research also shows that silencing of the *serotonin N-acetyltransferase 1* (*GhSNAT1*) and *caffeic acid O-methyltransferase* (*GhCOMT*) melatonin biosynthesis genes compromise cotton resistance, with reducing lignin and gossypol levels after *V. dahliae* inoculation ^[18]^.

A key player of pathogen resistance is the plant cell wall, which acts as a barrier to biotic and abiotic agents, and consists mainly of polysaccharides such as cellulose, hemicelluloses, lignin, and pectin. Degradation of plant cell wall compounds is a complex process involving the synergistic action of a large number of extracellular enzymes ^[19]^. Among all the components in the plant cell wall, pectin is arguably most complex and heterogeneous ^[20]^. Unfortunately, phytopathogenic fungi secrete pectinases that break down the middle lamella in plants, allowing for the extraction of nutrients from plant tissues and the insertion of fungal hyphae ^[21]^. The first cell wall degrading enzyme (CWDE) shown to be required for complete virulence was a pectate lyase published in 1985 from the bacterium *Erwinia chrysanthemi* ^[22]^. Since then, an increasing amount of evidence for the function of fungal pectinase has reported and these enzymes have regained the attention of plant pathologists. A pectate lyase (*VdPEL1*) exhibits pectin hydrolytic activity, which can induce drastic cell death in several plants ^[23]^. The gene *PG1*, which encodes for an endo-polygalacturonase, has been well studied in another plant vascular wilt pathogen, *Fusarium oxysporum,* and it plays a potential role in various aspects of fungal pathogenicity ^[24]^. *PG1* is also known to play an important role in depolymerizing homogalacturonan, a major component of the plant cell wall ^[25]^. *Pectin methylesterase (PME)* is an enzyme involved in the first step of the pathway that produces pectate from pectin. PME genes are present in both fungi and plants, and presumably play a role in cell wall remodeling. Interestingly, previous research has shown that there is a greater transcript abundance of fiber-specific PMEs and a higher total PME enzyme activity in *G. barbadense* (Gb) than in *G. hirsutum* (Gh) fibers, particularly during late fiber elongation ^[26]^. However, the function of these genes remains unknown in *V. dahliae*.

Mitogen-activated protein kinase (MAPK) signaling pathways have been shown to play key roles in infectious structure formation and in invasive growth of phytopathogenic fungi attacking the aerial parts of the plant, including the cereal leaf pathogens *Magnaporthe grisea* and *Cochliobolus heterostrophus* ^[27, 28]^, the maize pathogen *Ustilago maydis* ^[29]^ and the broad host range necrotrophic *Botrytis cinereal* ^[30]^. Some MAPKs are also shown to control the virulence of *V. dahliae*, including *VdHOG1* ^[31]^ and *VdPbs2* ^[32]^, promoting the virulence by regulating the formation of microsclerotia as well as *Vayg1* ^[33]^. The transcription factor (TF) *STE12* was found to regulate the expression of a polygalacturonase-encoding gene whose activity is related to the degradation of plant cell wall ^[34]^. In *S. cerevisiae*, *Ste12p* regulates mating and invasive/pseudohyphal growth. In phytopathogenic fungi, *Ste12* knockout strains are impaired in the development of the highly melanized appressoria and reduced virulence is shown for *Cryphonectria parasitica*, *B. cinerea*, *A. brassicicola*, and *F. oxysporum* ^[35]^.

Host-induced gene silencing (HIGS) has emerged as a promising strategy for the improvement of plant resistance against pathogens by silencing genes that are essential for pathogens using host miRNAs. For this reason, attention has been shifted to more intensive study of the pathogenetic fungi to achieve an effective control strategy towards VW diseases. Thus, research on the growth and pathogenic capability of the fungi at molecular level has become a cutting-edge and pivotal component of research on this problem. The new approach is generating more interest in the scientific community as evidenced by a recent increase in the number of publications in this new field. *Chitin-binding lysin motif* (*LysM*) effectors, which are found in a wide range of plant and animal fungal species, contribute to the virulence of *VdLs.17* ^[36, 37]^. The catalytic subunit of *cAMP-dependent protein kinase A* (*PKA*) is also studied to confer virulence within *V. dahliae* and some other plant pathogens ^[38, 39]^. Both are potential targets for HIGS.

In this study, we searched for key pathogenicity-related genes of *V. dahliae* by combining comparative transcriptomics and gene expression clustering. There was no precedent cluster analysis known on gene expression connectivity in the studies of *V. dahliae*. By using the D strain *Vd_086* and ND strain *Vd_BP_2_*, we identified a set of pectinases that likely promote the stronger virulence in *Vd_086* and these genes are possibly regulated by TFs in the MAPK signaling pathway. This study provides an important basis for the subsequent studies on pathogenetic gene discovery as well as the revealing of their functions and interactions.

## Results

### Defoliating (D) *Vd_086* and non-defoliating (ND) *Vd_BP_2_*

Characteristic attributes of the defoliating (D) *Verticillium dahliae* strain *Vd_086* and non-defoliating (ND) strain *Vd_BP_2_* clearly showed the difference in their morphology during cultured on the Potato Dextrose Agar (PDA) dishes (Fig. 1C), as well as on pathogenic capability of infecting Upland cotton plants (Fig. 1B, D). *Vd_086* (D) caused whole plant defoliation at 21 days post inoculation (dpi) with 10^7^ conidia/mL fungal spore suspension while most of the leaves of cotton plants inoculated by *Vd_BP_2_* (ND) still stayed green at the same time point (Fig. 1B). The phenotypic difference also was very apparent to the extent that browning of vascular bundles, which was more severe in *Vd_086* infected plants than *Vd_BP_2_* infected plants (Fig. 1A). Disease index ^[40]^ was utilized to measure the pathogenicity of the two *Vd* strains on cotton plants, demonstrating a significant difference from the control and from each other after the 13^th^ dpi (Fig. 1D). These data suggested that the D strain *Vd_086* has a stronger pathogenicity at the early invasion period. To learn what aspect of fungi caused the variation of early disease symptom, we quantified the growth rate of hyphae elongation and microsclerotia production during the entire medium culture period by measuring the length of diameters. Interestingly, it was observed that *Vd_086* is faster in hyphal growth but showed delayed black microsclerotia production (Fig. 1C). Although with the relatively less black microsclerotia, *Vd_086* showed a greater re-germination capability as evidenced by copious white mycelium passing through the top of the dish (Fig. 1C). The combined results from disease index and colony diameter measurements showed that the stronger pathogenicity in the early invasion phase of *Vd_086*, is possibly due to its rapid mycelial elongation and spore germination capability. On the other hand, the ND strain *Vd_BP_2_* caused milder pathogenic phenotype, but it would leave much black microsclerotia upon the death of infected cotton plants, which is also detrimental to subsequent cotton production (discuss below).

### RNA-Seq and genome mapping

To further reveal molecular mechanisms behind the stronger pathogenicity of the D *V. dahliae* strain through comparative analysis, RNA-seq was performed by using fungal samples collected during growth on PDA media. Samples from three phases of fungal growth, including hyphal growth (H stage), microsclerotia production (M stage), and spore germination (S stage), were selected in both D and ND *Vd* strains. With three replicates for each sample, there were 18 samples in all for RNA sequencing.

A total of 110.5 Gb clean sequence data were obtained with an average of 6.14 Gb for each cDNA sample. The minimum Q30 value was 88.49% with GC contents varied from 58.55% to 59.62%. For the joint analysis of all samples, we mapped the clean reads to a published *V. dahliae* strain *VdLs.17* genome that was downloaded from the Broad Institute ^[41]^. On average, 82.7% clean reads were mapped in which 81.8% were aligned to unique genomic locations (Table 1). Additionally, 703 new genes unmapped to the reference genome were discovered in which 295 were functionally annotated. Those genes were designated as *Verticillium_dahliae_newGene_001* to *Verticillium_dahliae_newGene_703*. With NR annotation, 96.17% mapped genes were classified to *Verticillium dahliae* database, but the rate was 34.01% in unmapped new genes. Most of new genes (44.56%) were classified as *Verticillium longisporum* genes, the other 11.56% were annotated to be involved in *Verticillium alfalfae* species (Fig. 2A). These data indicated that the genes in *Vd_086* and *Vd_BP_2_* not mapped to *VdLs.17* genome had a closer relationship with *Verticillium longisporum.* Among the total of 11,238 genes, 575 transcription regulators were annotated based on plantTFDB and animalTFDB database. Among them, there were 313 TFs, 134 transcription regulators, and 128 protein kinases (Fig. 2D). With further classification, most of the TFs were classified to C_2_H_2_ and Zn-clus families ^[42]^. Zn-clusters are usually specific to fungi as they have been detected in all fungal species analyzed so far ^[43]^. There was also a number of protein kinases annotated to be involved in CMGC (named after the initials of its subfamily members: CDK, cyclin-dependent kinase; MAPK, mitogen-activated protein kinase; GSK, glycogen synthase kinase and CLK and CDC-like kinase), CAMK (Ca^2+^/calmodulin-dependent protein kinase) and STE families. CMGC, a gene with some function is highly conserved across organisms. CaMK is a diverse group of protein kinases which activation is dependent on binding to Ca^2+^/calmodulin (CaM) complex. STE kinase consists of three main families that sequentially activate each other for the MAPK gene family.

**Fig. 2.**
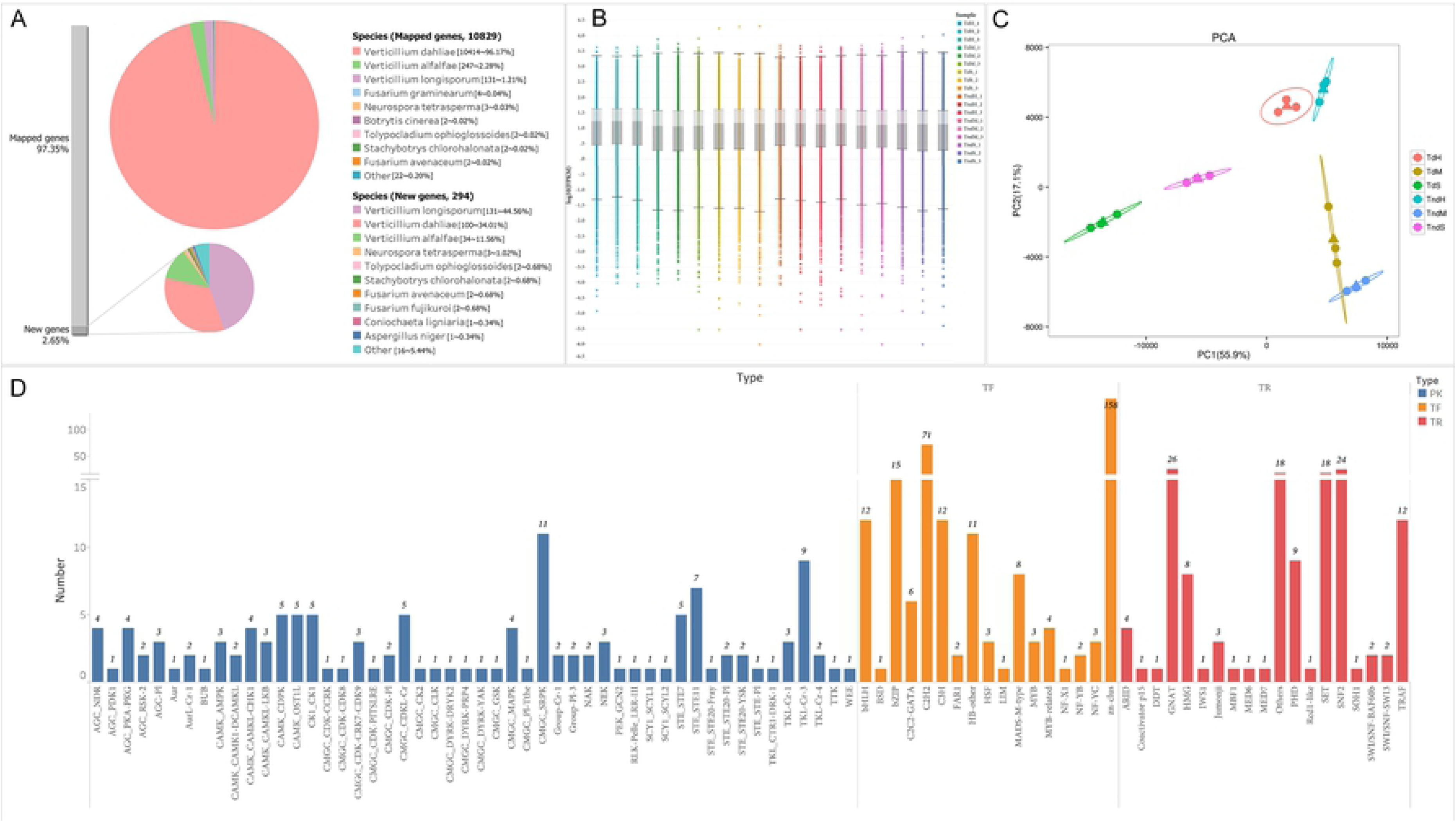
RNA-seq data display. (A) Species distribution of homologous genes mapped (large pie chart) and unmapped (small pie chart) to *VdLs.17* genome. (B) Box plot of all gene expression (fpkm) in 18 samples. (C) Principle component analysis (PCA) of all expressed genes. The abscissa and ordinate, respectively, represent the first and second principal components, and the contribution of each principal component is in parentheses. Different samples are distinguished by color, with three replicates for each group. The triangle as the center point represents the mean coordinate of each group, and the ellipse represents the confidence interval. (D) Distribution of predicted transcription factors (TFs). Different TF types are distinguished by color.

**Table 1.**
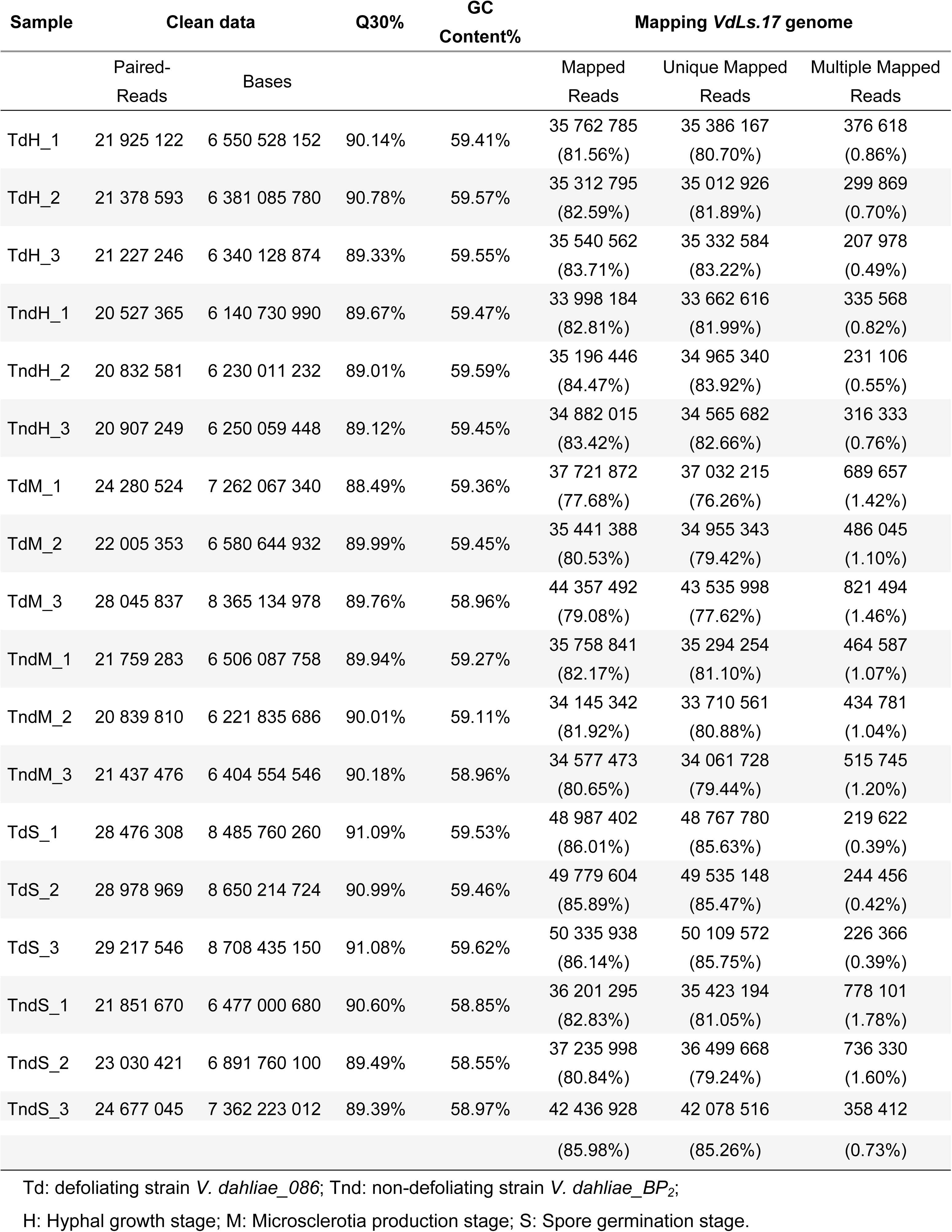
Raw data of RNA sequencing.

### Comparative analysis with differentially expressed genes (DEGs) between two pathotypes of *V. dahliae*

Box plot was drawn to display the distribution of gene expression (value of log_10_fpkm) along each sequenced sample (Fig. 2B). The gene expression distributed consistently among replicates but slightly different between samples in different groups. The first two axes of principal component analysis (PCA) of the gene expression data explained 55.9% and 17.1% variation, which also showed a great repeatability in each group of samples (Fig. 2C). There were 1,098, 1,123, and 820 DEGs identified at hyphal growth (H), microsclerotia production (M), and spore germination (S) phases, respectively. Among them, 480, 683, and 403 DEGs were unique at a single growth stage, and 348 DEGs were conserved at all three growth stages (Fig. 3B). Compared with the ND strain, the number of down-regulated genes in the D strain was about twice the up-regulated genes at every stage, indicating more active gene expression in the ND strain (Fig. 3A).

**Fig. 3.**
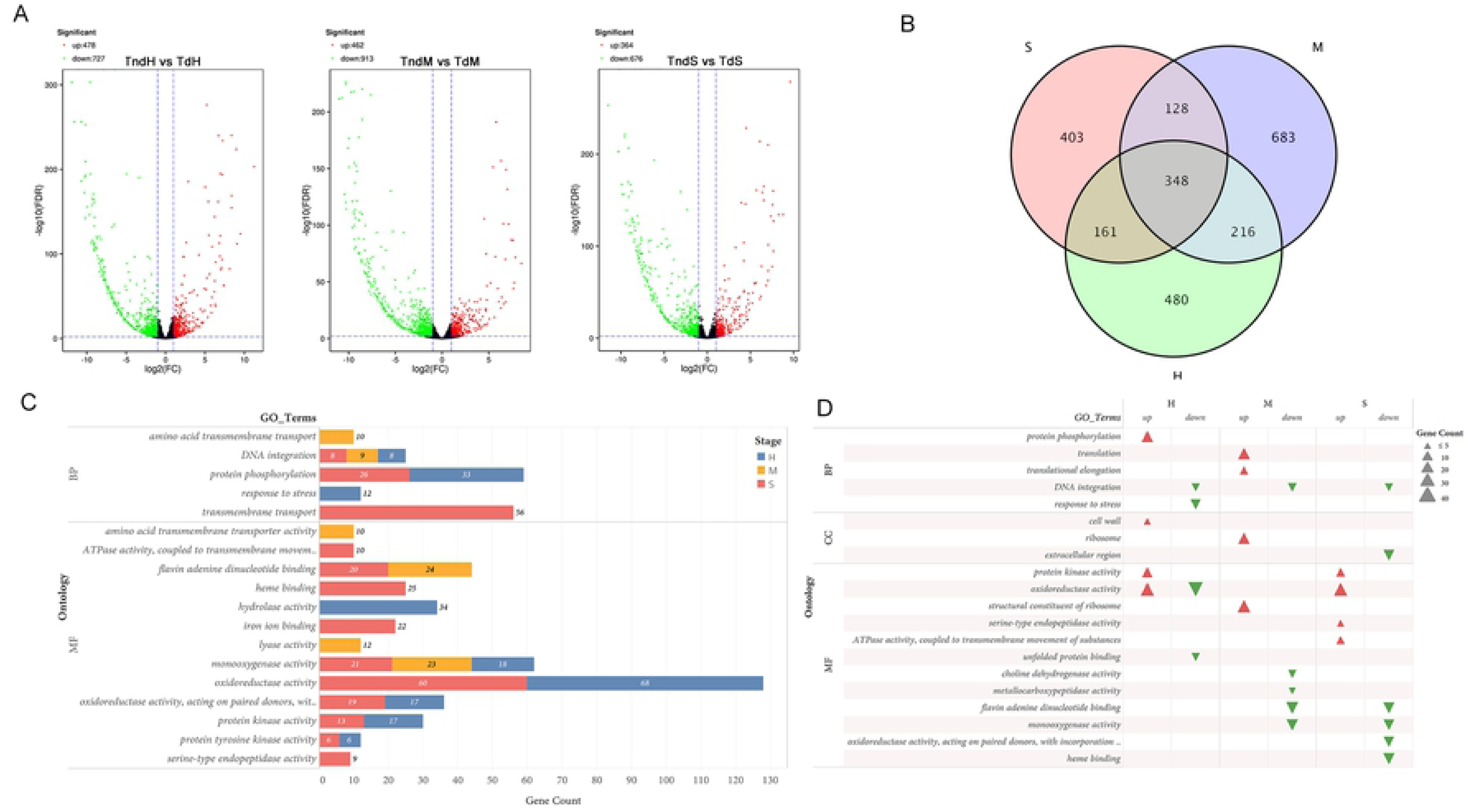
Identification and functional enrichment of differentially expressed genes (DEGs). (A) Volcano plots of DEGs within *Vd_086* compared to *Vd_BP_2_* at hyphae (H), microsclerotia (M) and spore germination (S) stages separately. (B) Venn plot of DEG numbers at three *V. dahliae* growing stages. (C) Gene Ontology (GO) enrichment. Terms with p < 0.005 are displayed of DEG sets. (D) GO enrichment. Terms with p < 0.005 of directionally expressed DEG sets.

Functional enrichment of differentially expressed genes (DEGs) between *Vd_BP_2_* (ND) and *Vd_086* (D) shed light into molecular mechanisms behind stronger pathogenicity and faster initial growth of the D strain. The functional enrichment was analyzed using the three DEG datasets (H, M, and S) by ClusterProfiler in R with all 6,255 GO-annotated genes as the background. Those DEGs significantly enriched at the p-value (< 0.005) were displayed in (Fig. 3C). Major differences between the two strains at H stage were observed in the expression of several hydrolases, oxidoreductases, and protein kinases. Interestingly, DEGs at the S stage were also significantly enriched in oxidoreductase activity and protein kinase-related activities. Since spore germination and mycelial elongation belong to the initial phases of development of *V. dahliae,* which was crucial for the pathogenic fungus to infect host plants, different activities of hydrolases, oxidoreductases, and protein kinases were the possible reason for the earlier infection of the D strain.

To further define the gene expression directionality in the D strain than to the ND strain, the up- and down-regulated gene sets of the DEGs at each stage were functionally enriched (Fig. 3D). During the early developments (H and S stages), all the DEGs significantly enriched in the protein kinase activity showed higher expression in the D strain, showing a relationship between protein kinases and *Vd* early pathogenicity. These gene candidates were listed in (S3.a Table). On the other hand, the DEGs enriched in oxidoreductase activity displayed a higher expression in the D strain on S stage, but a mixed expression on H stage. Additionally, the D-up genes (genes with an increased expression in the D strain) were enriched in genes involved in “protein phosphorylation” and “cell wall” structures, implying a possible relevance of these processes towards the fast development and rapid plant infection of the D strain. Of interest was the enrichment in “response to stress” and “unfolded protein binding” processes of D-down genes (genes with decreased expression in the D train) on H stage. This might explain the D strain’s weaker production capability of microsclerotia which is known as stress response structure. At the M stage, genes for translation and ribosome-related process were highly expressed in the D strain. These activities might inhibit or delay the formation of microsclerotia. Genes for DNA integration were expressed conservatively low in the D strain at all three growth stages, indicating a weaker capability of foreign genetic materials that is inserted in the D strain.

Twelve KEGG pathways were significantly enriched using the DEG sets with the q-value of FDR smaller than 0.05 (S1.a Fig). The metabolic pathways of several amino acids like arginine, proline, glycine, serine, and threonine were possibly related to the *V. dahliae* hyphal elongation. Also, the significant enrichment of “<u>p</u>entose and <u>g</u>lucuronate interconversions” (PGI) and “phenylalanine metabolism” for gene sets at both H and S stages presented a likely correlation of these genes towards the initial development of the D strain. From the enrichment of directional expression gene sets, the PGI genes were very active in the D strain at the early development stages (H and S) (S1.b Fig; S3.b Table). In addition to molecular functions such as “amino acid transmembrane transport”, DEGs at M stage were also enriched in the coherent processes of biosynthesis of antibiotics, carbohydrate metabolism, and ribosome. Only the ribosome and amino acid biogenesis genes showed active expression in the D strain while some fatty acid degradation and tryptophan metabolism genes were active in the ND strain. This might explain the reason why microsclerotia in *Vd_BP_2_* was much darker in color as tryptophan was verified to be a non-canonical melanin precursor ^[44]^.

### Comparative analysis with weighted gene expression clustering

Identification and annotation of DEG sets for mining key genes is a common approach to comparative genomics. However, as alterations of numerous transcripts occurred in multiple terms and pathways, it is challenging to pinpoint which specific DEG plays a causal role in the more severe pathogenicity of the D *Vd* strain. It is also difficult to identify key regulatory genes. Thus, weighted gene co-expression network analysis (WGCNA) was performed to further explore the specific molecular mechanisms and key genes that affected three stages of *V. dahliae* growth within two pathotypes. An fpkm data set with 6,875 genes that passed the filtering criteria (parameters see Methods) was utilized as input data. There were no outlier samples needed to be removed by offending genes of sample screening and the soft power was set at 22 because this was the lowest power needed to reach approximately scale-free topology (R^2^=0.865) (S2 Fig). With the minimum merging height changed from 0.15 to 0.25 in dendrogram cutting, the number of co-expression modules reduced from 30 to 23 (Fig. 4A).

**Fig. 4.**
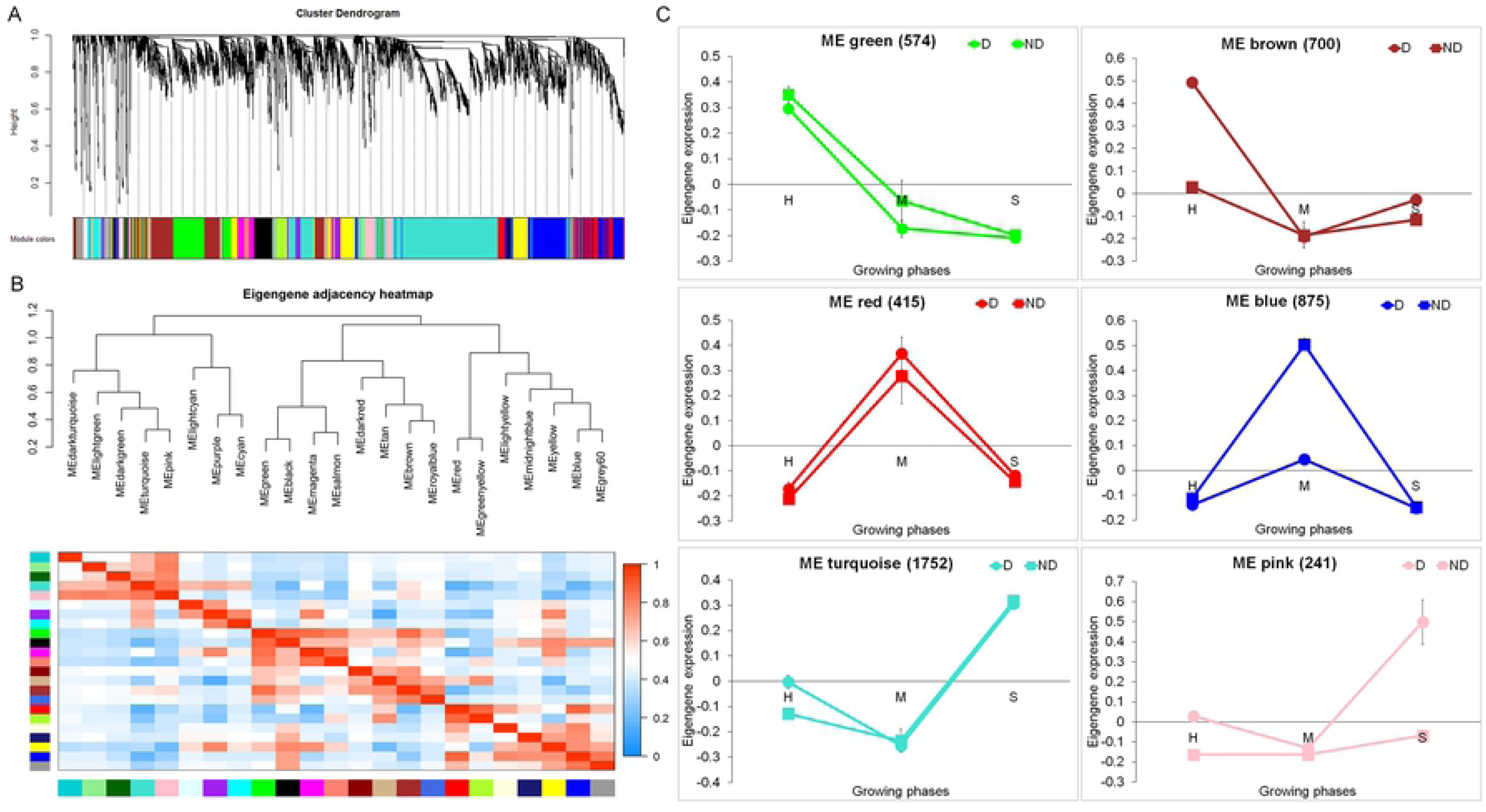
Weighted gene correlation network analysis (WGCNA). (A) Hierarchical clustering of the topological overlap matrix for *V. dahliae* expression data. (B) Hierarchical clustering of the 23 detected modules. The heatmap shows the pairwise Pearson correlations. (C) Six attributes matching modules of co-expressed genes during development of *V. dahliae*. The RNA-seq expression data set was subjected to WGCNA to detect modules of co-expressed genes. Each graph shows the expression of the module eigengene, which can be considered as the representative gene of the respective co-expression module. The vertical axes indicate log2 expression values relative to the mean expression across all stages. The horizontal axes indicate the stages H, M, and S. Error bars represent standard deviations (SD) of three biological replicates. The modules are named according to their color, and the number of genes residing in each module is given in parentheses.

Modules associated with the stronger pathogenicity and more rapid early-development of the D strain vs. the ND strain are shown in (Fig. 4C). Among all 23 clustered gene modules, the eigengene expression trends of six gene modules showed patterns that coincide with the attributes of the *V. dahliae* growth (Fig. 4C). Green, red, and turquoise module eigengenes shared similar expression patterns between two *Vd* strains with a single peak at H, M, and S stages, respectively (Fig. 4C). These data indicated that the gene expression in these modules is conserved between the two strains and likely contributes to important functions in the normal growth of both fungal strains. However, genes in brown and pink had a higher expression in the D strain than the ND strain at either H or S stage that correlated with the early fungal growth and host infection in *Vd_086*. The blue module eigengenes displayed an opposite trend with a higher expression in the ND strain at M stage, which may be related to an earlier production of darker microsclerotia in *Vd_BP_2_*.

To better examine the details of gene expression during the pathogenesis process, we divided the six WGCNA co-expression modules into three comparison groups based on each growth phase. In view of eigengene expression trends and module co-relationships (Fig. 4B, C), modules green (genes with a common pattern of expression in both *Vd* strains in the hyphal stage) and brown (genes with a higher level of expression in the faster growth *Vd_086* (D) strain at the hyphal stage) were paired for comparing enriched functional gene categories (GO terms were significant with p-value < 0.005) (Fig. 5A). Transmembrane transport (green) was a functional category that was enriched in in both *Vd* strains during the elongation stage. Genes with a higher level of expression in *Vd_086* (D), in the brown module, were enriched in the categories of “proteolysis” and “carbohydrate metabolic processes”, suggesting that these may be important in the more rapid hyphal elongation in this strain (discussed below) (Fig. 5A: green & brown). Subsequently, “biogenesis of ribosome” and “ATP-dependent helicase” processes were activated during microsclerotia formation. Some gene activities appeared to be very active during spore germination when the environment was suitable for initiating a new life cycle of *Vd* infection (Fig. 5A: turquoise). However, it was not known what caused the faster spore germination in the D strain as the pink module genes were not significantly enriched with our threshold for GO enrichment - this may be due to the small number of genes (241) in pink module.

**Fig. 5.**
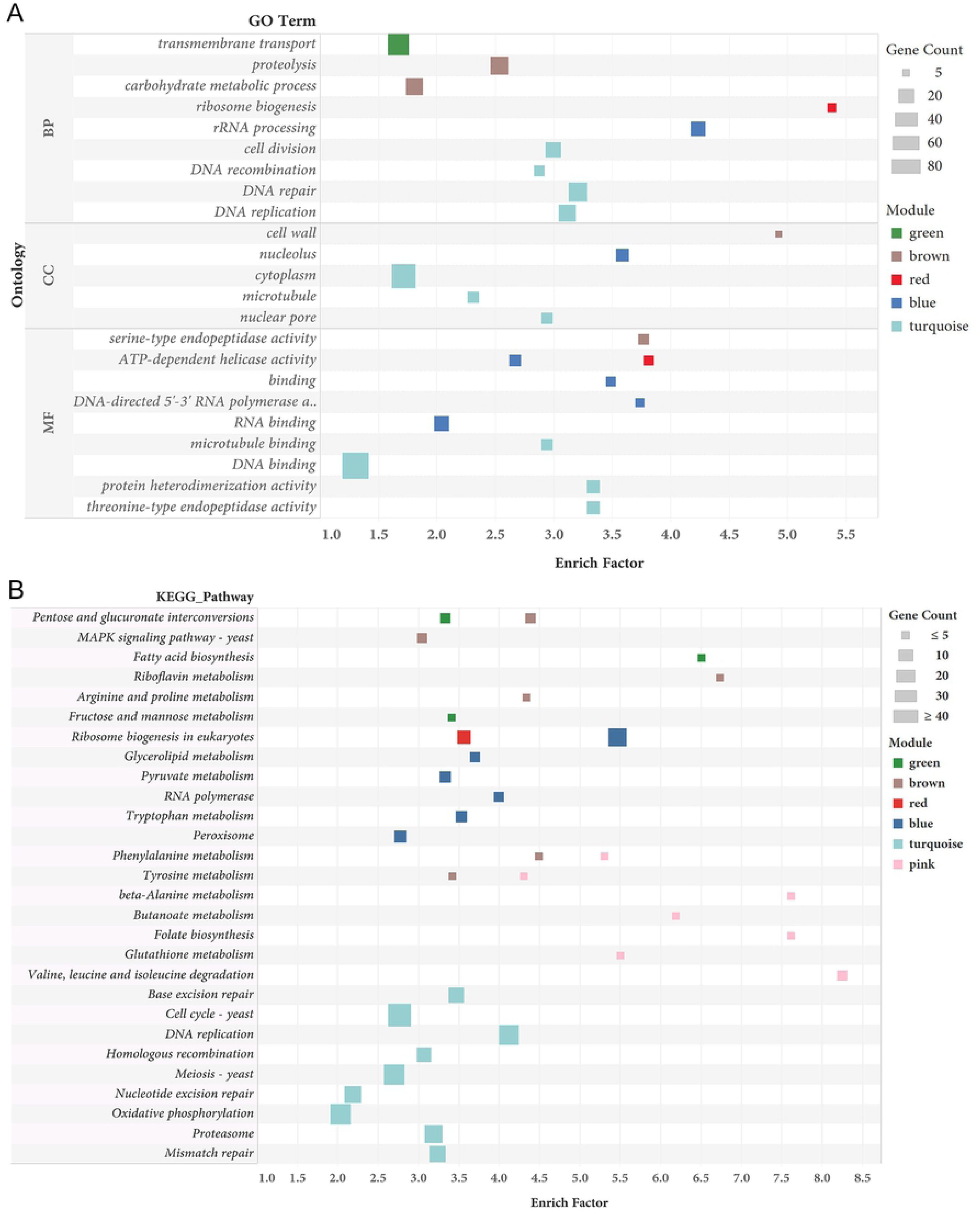
Functional enrichment of six module gene sets. (A) GO term enrichment with p < 0.005. (B) KEGG pathway enrichment with q(FDR) < 0.25.

KEGG enrichment was performed to further understand the key pathways involved in the *Vd* growth. Pathways significant with q-value (FDR) < 0.25 were selected and displayed in Fig. 5B. Similar to the GO result, metabolism of some carbohydrates like pentose, fructose, mannose, and fatty acid was responsible for the normal hyphal growth of *Vd* (Fig. 5B: green). Surprisingly, genes in the brown module (represented the faster mycelium elongation), which were enriched in the PGI pathway (gene count = 6). In addition, brown genes were involved in amino acid metabolism and in MAPK signaling pathway, a widely accepted pathogenetic process in plant pathogens (Fig. 5B: brown). After that, ribosome biogenesis was likely activated during the formation of microsclerotia (Fig. 5B: red) and the darker phenotype of the ND strain was possibly conferred by more active RNA associated proteins and some melanin related amino acids like tryptophan (Fig. 5B: blue). The metabolism of phenylalanine and tyrosine was identified by pink module (representing the faster spore germination) genes, indicating a possible relationship towards the difference of the initial development between two *Vd* strains.

### Pectin degrading capability of defoliating (D) *Vd_086*

The pathway of pentose and glucuronate interconversions (PGI, ko00040 in KEGG database) caught our attention because of the significant enrichment by DEG and gene clustering modules associated with *V. dahliae* hyphal growth, that involved pectinases. As a major component of plant cell wall, the structure of pectin is a chain mainly consisted of poly (1,4-α-D, galacturonide), whose degradation provides carbohydrate raw materials for the entire PGI pathway (Fig. 6A). Therefore, we suggest that *V. dahliae* secretes pectinases to degrade cell wall of the host plant in order to obtain the source of carbohydrates needed for fungal metabolism and growth, enhancing its infection and pathogenicity along the way, especially in the D strain.

**Fig. 6.**
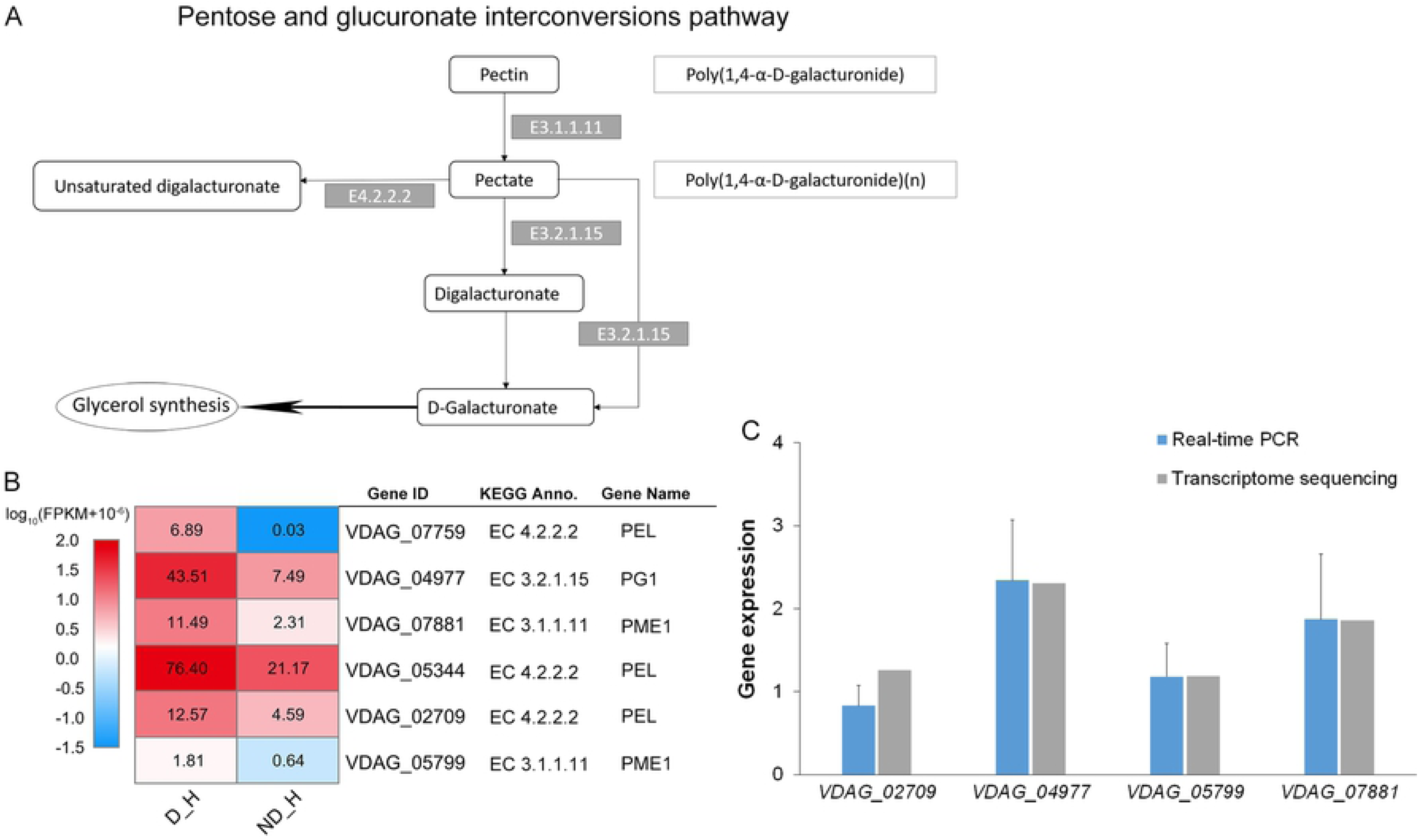
Gene expression of pectin degrading enzymes. (A) Overview of pentose and glucuronate interconversions (PGI) pathway. (B) Expression profile of different expressed pectin degrading enzymes in the PGI pathway during hyphal stage. D_H, defoliating strain *Vd_086* in hyphal stage. ND_H, non-defoliating strain *Vd_BP_2_* in hyphal stage. The expression levels are indicated by the heatmap, estimated using log10 (FPKM + 10^-6^) for each transcript. The data on the grids of heatmap was gene FPKM value. (C) Comparison of relative gene expression between RNA-seq and qRT-PCR. Gray bars represent log2 (fold change) of gene expression in *Vd_086* comparing to *Vd_BP_2_*. Blue bars represent a mean value of 2^-ΔΔCt^. Error bars represent standard deviation (SD) of three biological replicates.

To verify whether the faster mycelial elongation and earlier pathogenicity of the D strain were resulted from certain degrading enzymes in PGI, we examined all the PGI genes for differential expression (DE) analysis. Twenty-eight genes remained after the removal of low expressed genes (fpkm < 1.0 in all samples) (Table 2). Seven of them showed higher expression (log2fold change > 1) in the D strain than the ND at H stage (genes with * in Table 2). Surprisingly, six of these genes conferred core pectin degrading enzymes at the top part of PGI pathway. They were annotated via Swiss Protein Database as *PG1* (*VDAG_04977*), *PEL* (*VDAG_02709, VDAG_07759, VDAG_05344*), and *PME1* (*VDAG_07881, VDAG_05799*) (Fig. 6B). Based on the previous studies, the building block of pectin is poly(1,4-α-D, galacturonide), the glycosidic bond can be cleaved by hydrolysis (using endo-polygalacturonases (*PGs*) [EC 3.2.1.15], exo-polygalacturonases [EC 3.2.1.67], and by a nonhydrolytic reaction called β-elimination, which uses pectin lyases [EC 4.2.2.10], pectin methylesterases (*PMEs*) [EC 3.1.1.11], and pectate lyases (*PELs*) [EC 4.2.2.2] ^[35, 45]^ (Fig. 6A). Thus, among the six highly expressed pectin degrading enzymes, four of them were verified with qRT-PCR which showed a high correlation to the RNA-seq data (Fig. 6C). These results demonstrated that the rapid mycelial development and pathogenic capability of the D *V. dahliae* strain were associated with strong pectin degradation activity.

**Table 2.** Twenty-eight genes involved in the PGI pathway after removing low expressed genes.

| Gene ID | Module | Gene expression (FPKM) |  | EC_ID | Gene Name | Pfam annotation |
| --- | --- | --- | --- | --- | --- | --- |
|  |  | D_H | ND_H |  |  |  |
| VDAG_07759* | - | 6.89 | 0.03 | E4.2.2.2 | *PEL* | Pectate lyase |
| VDAG_04977* | brown | 43.51 | 7.49 | E3.2.1.15 | *PG1* | Glycosyl hydrolases family 28 |
| VDAG_07881* | - | 11.49 | 2.31 | E3.1.1.11 | *pme1* | Pectinesterase |
| VDAG_05344* | - | 76.4 | 21.17 | E4.2.2.2 | - | Pectate lyase |
| VDAG_00549* | brown | 42.08 | 13.45 | E1.1.1.14 | - | Alcohol dehydrogenase GroES-like domain;<br>Zinc-binding dehydrogenase |
| VDAG_02709* | brown | 12.57 | 4.59 | E4.2.2.2 | PEL | Pectate lyase; Right handed beta helix<br>region |
| VDAG_05799* | - | 1.81 | 0.64 | E3.1.1.11 | pme1 | Pectinesterase |
| VDAG_05087 | - | 4.55 | 2.48 | E1.1.1.307 | PRD1 | Aldo/keto reductase family |
| VDAG_08773 | - | 173.03 | 116.65 | E5.1.3.1 | J0731 | Ribulose-phosphate 3 epimerase family |
| VDAG_08929 | green | 1513.32 | 1028.83 | E1.1.1.287 | ARD1 | Alcohol dehydrogenase GroES-like domain;<br>Zinc-binding dehydrogenase; Zinc-binding<br>dehydrogenase |
| VDAG_01781 | - | 6.05 | 4.36 | E3.2.1.67 | pgaX-2 | Glycosyl hydrolases family 28 |
| VDAG_02126 | - | 228.77 | 175.66 | E1.1.1.22 | FLO11 | UDP-glucose/GDP-mannose<br>dehydrogenase family NAD binding domain;<br>central domain; UDP binding domain |
| VDAG_10081 | - | 34.38 | 27.61 | E42.1.146 | lgd1 | Enolase C-terminal domain-like; Mandelate<br>racemase / muconate lactonizing enzyme<br>C-terminal domain |
| VDAG_04953 | brown | 64.99 | 53.07 | E1.1.1.127 | DHG2 | Enoyl - (Acyl carrier protein) reductase;<br>short chain dehydrogenase |
| VDAG_05992 | - | 7.17 | 5.91 | E3.2.1.67 | pgxB | Glycosyl hydrolases family 28 |
| VDAG_03025 | - | 236.97 | 211.89 | E2.7.7.9 | fuy1 | UTP--glucose-1-phosphate<br>uridylyltransferase |
| VDAG_06079 | - | 41.45 | 38.05 | E1.1.1.14 | ard-1 | Zinc-binding dehydrogenase |
| VDAG_00768 | - | 10.49 | 10.12 | E3.2.1.67 | pgxB | Glycosyl hydrolases family 28 |
| VDAG_01073 | green | 244.95 | 227.42 | E1.1.1.307 | - | - |
| VDAG_03683 | - | 235.55 | 219.66 | E1.1.1.372 | gld1 | Aldo/keto reductase family |
| VDAG_04952 | brown | 27.89 | 26.71 | E1.1.1.14 | xdhA | Alcohol dehydrogenase GroES-like domain;<br>Zinc-binding dehydrogenase |
| VDAG_08495 | green | 37.27 | 41.31 | GaaA | - | Oxidoreductase family NAD-binding<br>Rossmann fold |
| VDAG_05815 | - | 1.17 | 1.34 | E2.7.1.17 | - | - |
| VDAG_07810 | - | 23.08 | 34.75 | E1.1.1.372 | gld1 | Aldo/keto reductase family |
| VDAG_08496 | - | 2.14 | 3.74 | E.1.2.54 | - | - |
| VDAG_05402 | - | 4.16 | 7.78 | E4.2.2.2 | PEL | Pectate lyase |
| V DAG_06181 | green | 50.55 | 101 | E1.1.1.9 | - | - |
| V DAG_00411 | - | 0.02 | 9.78 | E1.1.1.307 | - | - |

Gene IDs with \* represent significantly higher expressed genes in *Vd\_086* (D) strain; D\_H: defoliating strain *Vd\_086* in hyphal growth stage; ND\_H: non-defoliating strain *Vd\_BP<sub>2</sub>* in hyphal stage.

### Function of mitogen-activated protein kinase (MAPK) signaling transcription factors

Since the earlier pathogenicity and elongation capability of the D strain were closely related to active pectin degrading enzymes, the transcription factors (TFs) regulating this process were explored. The possible TFs with 2 kb nucleotides before a gene’s start codon were predicted as gene promoters in fungal database by PROMO ^[61]^. To further examine the interaction between genes and TFs at the hyphal development stage, the amino acid sequences of the brown and green module genes were loaded into the STRING database (v11.0) ^[65]^ to generate the protein-protein interaction (PPI) network based on species of *Verticillium dahliae* and visualized with Cytoscape (Fig. 7A).

**Fig. 7.**
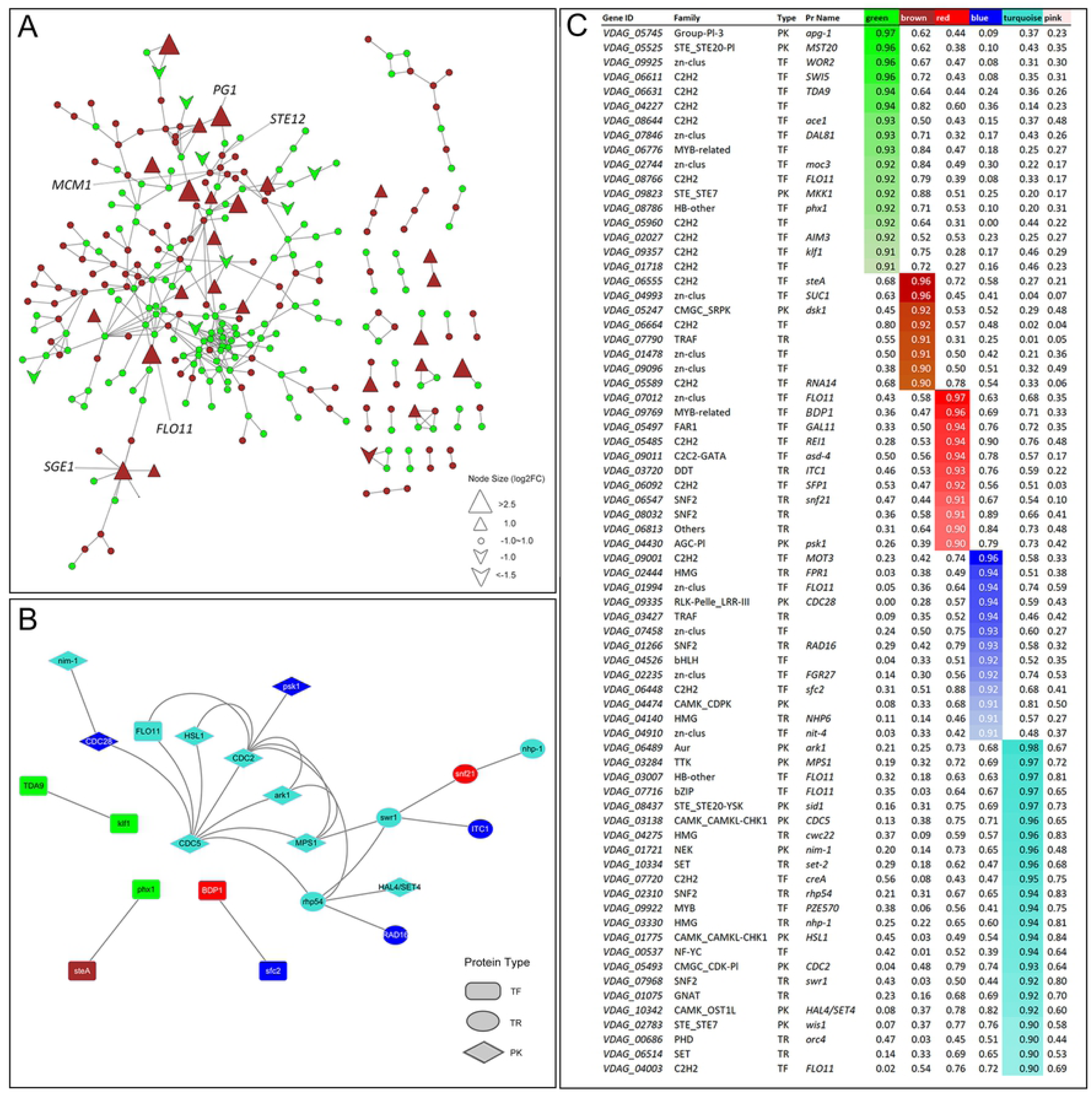
Hub genes and PPI network. (A) Protein to protein interaction (PPI) network of green and brown module genes. The shape of nodes represent the gene expression mode in *Vd_086* comparing to *Vd_BP_2_*. Node size of triangle and V shape represents the gene expression level estimated using log2 (fold change). Genes are colored according to their modular membership. Some key genes are labeled with their respective names. (B) Network of hub transcription factors belonging to the respective modules in which hub genes (kME > 0.9) reside. Proteins are colored according to their modular membership. TF: Transcription factors; PK: Protein Kinases; TR: Transcription regulators. (C) Connectivity of transcription factors (TFs) to the six modules. Depicted are all TFs having an intramodular connectivity of greater than 0.9 to any of other modules. Color intensity indicates connectivity strength.

Through the PPI network (290 nodes, 401 edges) of the brown-green module, it was shown that the brown module proteins were very active at the mycelial development stage of the D strain with a large number of genes that were significantly higher expressed (Fig. 7A). Based on PROMO, three TFs (FACB, GA-BF, RC2) were predicted to regulate those six DEGs involved in pectin degrading process but none of them was identified in the brown module. A protein *STE12* was identified as a potential regulator of the most highly-expressed pectinase *PG1 (VDAG_04977)* by the presence of sequence of TGTTTCAT with its promoter. As a target of the pheromone-induced signal in the mitogen-activated protein kinase (MAPK) signaling pathway (ko04011 in KEGG database), *STE12* acts with another pheromone receptor TF *MCM1* to express optimal transcriptional activity for the mating of fungal mycelia ^[46]^. *MCM1* is an intra-modular TF within the brown module and, based on PPI, is associated with many proteins enriched in brown and green modules (Fig. 7A). Furthermore, *STE12* was also required for expression of *FLO11* which was manifested to be an hub intra-modular green-module protein to promote the fungal filamentation ^[47]^.

Highly-connected “hub genes” were identified by kME value. With kME > 0.9 set as the parameter for intra-modular hub gene selection, 72 TFs were identified as possible hub regulators within five modules (Fig. 7C) (none TFs in pink module met the threshold). The PPI network of all hub TFs were displayed in Fig. 7B. All these proteins were shown to regulate a number of genes in the module they were located. Several hub protein kinases were present in turquoise module indicated a relatively active protein modification process at spore germination stage of both D and ND strains. In addition to the hub TFs of green module, *FLO11* was also the core regulator of another three modules of red, blue, and turquoise. These data indicated that *FLO11* may be a crucial TF to regulate the entire growth and development of *V. dahliae*. In addition to *STE12*, *MCM1*, *FLO11* which involved in MAPK signaling, TFs like *MST20* (green), *MKK1* (green), *steA* (brown), *CDC28* (blue), *CDC2* (turquoise), and *wis1* (turquoise) were all involved in the MAPK signaling and acted as hub regulators in their modules (Fig. 7C).

## Discussion

The assessment of *V. dahliae* shows that the D strain exhibits much stronger initial pathogenicity during plant infection. On the other hand, the ND strain has an apparent advantage on microsclerotia production. Based on previous studies, microsclerotia exist as a resting structure of filamentous fungi for protecting themselves from severe environments ^[48]^. With the mycelium elongating, the nutrition was insufficient for further hyphal growth on medium and inside plant host. At this point, the phytopathogenic fungi produce a melanized, multicellular survival structure known as microsclerotia which is critical for long-term survival in soil. Some studies showed that the capability of microsclerotia production was responsible for the pathogenicity of *V. dahliae* ^[33]^, but an opposite result is shown in our research in that the ND strain with relative weak pathogenicity displayed a stronger black microsclerotia production capability. This implies that the prevention and control of both D and ND pathogens are equally important, because the ND pathogens likely leave more microsclerotia behind in the disease field, which is a serious problem to subsequent years. Thus, there is a considerable amount of work to identify functional genes during the growth pathogenesis of both D and ND *V. dahliae*.

Identification of differentially expressed gene (DEG) sets and their functional annotation is an effective strategy used in comparative transcriptome analysis. But it is not accurate enough to identify specific pathogenicity genes due to the large number of genes inside DEG sets. Recently, weighted gene co-expression network analysis (WGCNA) becomes a widely used bioinformatic tool that incorporates hierarchical clustering to identify modules with different tree-cutting thresholds through summarizing each module by the module eigengene or intramodular hub genes ^[49]^. Generally, with the combination of sample gene expression matrix and trait data matrix, phenotype-related gene clusters can be identified. But in the present study, it was difficult to quantify the whole-stage growth parameter of *V. dahliae*. Fortunately, the scaled module eigengene expression value was representative of the expression trends of specific gene modules. With the visualization of line graphs (Fig. 4C), modules matching the attributes of D and ND *V. dahliae* were clearly recognized. Followed by the DEG analysis, module gene functional annotation, and intramodular DEG recognition, the pentose and glucuronate interconversion (PGI) pathway (ko00040) was identified to be important in early fungal growth. A group of pectin degrading enzymes were identified as upregulated in the D strain, providing a plausible explanation for the faster mycelium elongation and stronger initial pathogenicity of the D strain.

*V. dahliae* enters the host plant through the wounded roots (usually caused by nematodes in soil), then colonizes the vascular bundles through the elongation of the mycelium. It penetrates the plant cell wall in this process by the secretion of many cell wall-degrading enzymes (CWDEs) ^[50]^. In filamentous fungi, these plant CWDEs provide them with the means to obtain energy and nutrients from plant cell wall biopolymers and are also believed to contribute to pathogenesis at the cellular level by breaking the cell barrier ^[51]^. Actually, all phytopathogenic fungi require these enzymes at invasion stages ^[52]^. The first CWDE shown to be required for complete virulence was a pectate lyase published in 1985 ^[22]^. Since then, additional evidence for the function of fungal pectinase has been reported ^[22–24]^ and these enzymes have regained the attention of plant biologists.

The main chain of pectin is composed of hairy and smooth regions. The hairy region is formed by its significantly sized side-chains. The structural differences between these two regions have implications for enzymes necessary to degrade these regions, with the hairy regions needing several additional accessory enzymes. In this study, six pectin degradation-related enzymes involved in the PGI pathway were identified. They were correlated to the stronger pathogenicity of the D strain and were annotated to be [EC 3.1.1.11], [EC 3.2.1.15] and [EC 4.2.2.2]. Four identified enzymes were shown by qRT-PCR to be highly expressed in the D strain, and the gene expression levels were consistent with the transcriptome sequencing data. Our results suggested that stronger early pathogenicity of the D strain is developed through enhanced plant cell wall pectin degradability. However, the specific mechanism of these pectinases awaits further experimental verification. Moreover, the correlation of intra-modular transcription factors (TFs) and genes can be predicted by the construction of gene network. With the TF prediction on PROMO, it was apparent that the mitogen-activated protein kinase (MAPK) signaling TFs were crucial regulators for *V. dahliae* development, in addition to the three TFs (FACB, GA-BF, RC2) identified by all seven pectin degrading enzymes. As the MAPK signaling associated genes are widely accepted for their involvement in the virulence of *V. dahliae*, we propose that the virulence provided by MAPKs may derive from their regulation of functional cell-wall degradation associated enzymes and genes. The regulation pathway in which the MAPKs are acting would require further experimental investigation.

In recent years, efforts have been made in discovering the molecular mechanisms underlying the interactions between phytopathogenic fungus *V. dahliae* and plants of *Arabidopsis*, tomato, potato, and cotton. Omics analysis, especially comparative analysis, is getting more extensively used in the identification of key genes. In this study, a new concept is employed for functional gene discovery through the combination of comparative transcriptomic analysis and weighted gene correlation network analysis (WGCNA). The specific functions of the genes implicated in this analysis, and their roles in the process of pathogenetic infection, remain to be confirmed with further experimentation. In the future, gene silencing in filamentous fungi with homologous recombination and CRISPR/Cas9 gene editing may be used to examine the fungal growth and pathogenicity through the construction of mutants that are absent of single genes. The microRNAs that are capable of regulating or inhibiting those functional genes may be tested by host induced gene silencing (HIGS) verification in cotton or other host plants. Those microRNAs, proven to successfully inhibit the expression of pathogenic genes within the host plants, may be utilized in plant molecular breeding programs. A new framework for functional studies of candidate genes will lead to better strategies to control the cotton VW disease.

## Materials and Methods

### Fungal strains, plant materials, and culture conditions

#### Sources of fungal strains and cotton plants

The defoliating (D) *Verticillium dahliae* strain *Vd_086* was isolated from Upland cotton plants in General Mongolian Autonomous Prefecture, Xinjiang, China ^[53]^. The non-defoliating (ND) strain *Vd_BP_2_* was provided by Institute of Plant Protection, Jiangsu Academy of Agricultural Sciences, Nanjing, China. Both *V. dahliae* strains were maintained in Cotton Genetics and Molecular Breeding Laboratory in College of Agriculture and Biotechnology at Zhejiang University. VW disease index was assessed using a susceptible cotton variety EJing-1 (EJ-1) that was provided by Plant Protection Institute, Hebei Academy of Agricultural and Forestry Sciences, China.

#### Fungal culture, attribute measurement and virulence assessment

*Verticillium dahliae* strains were grown on the Potato Dextrose Agar (PDA) medium at 25°C. Diameters of mycelial and dark microsclerotia were measured to assess the growth rate of *V. dahliae*. To collect conidia, mycelial plugs were cultured in Czapek-Dox liquid medium (30 g/L Sucrose, 3 g/L NaNO_3_, 0.5 g/L M gSO_4_-7H_2_O, 0.5 g/L KCl, 100 mg/L FeSO_4_-7H_2_O, 1 g/L K_2_HPO_4_, pH 7.2) with shaking at 150 rpm at 25°C in dark for 3 - 5 days. After centrifugation at 4,000 rpm for 10 min, followed by the resuspension in sterile distilled water, conidia at a concentration of 10^7^ conidia/mL were used for fungal inoculation ^[54]^. Each *Vd* treated cotton plant was inoculated with 15 mL spore suspension at two true leaves stage with root wounded by a long-handled spoon. The 5-grade disease index was used to measure the infection results as follows: 0 (no symptoms), 1 (1 - 2 cotyledons wilted or defoliated), 2 (1 - 2 true leaves wilted or defoliated), 3 (3 - 4 true leaves wilted or defoliated), and 4 (whole plant leaves wilted or defoliated). The disease index was first calculated at the 7^th^ day after inoculation (dpi) as 100 × [∑(number of plants × disease grade)] / [(total number of plants) × (maximal disease grade)] ^[40]^.

### RNA-Seq

#### Sampling of fungal tissues

Mycelial plugs were cut by 5-cm hole puncher on the edge of *V. dahliae* colonies grown on the PDA medium for 7 days. Each 5-cm mycelial plug was put on the middle of a new PDA plate overlaid with a cellophane layer for following samples collection ^[55]^. The hyphal stage (H stage) samples were collected from 3 d, 4 d, 5 d PDA plates that were incubated at 25°C in the light. The microsclerotia stage (M stage) samples were collected from 8 d, 10 d, 12 d PDA plates at 25°C in the dark. The 12-day dark cultured dishes were washed by suitable sterile water, and then the acquired spore suspension was coated on cellophane covering the PDA plates. Spore germination (S stage) samples were collected after 10 h and 20 h dark culture under 25°C. These H, M, S stage samples with three replicates of each were used for RNA-seq and further expression identification.

#### RNA extraction and library preparation

Total RNA was extracted using the RNAprep Pure Plant Kit (Tiangen, Beijing, China) according to the manufacturer’s instructions. RNA concentration was measured using NanoDrop 2000 (Thermo Scientific). RNA integrity was assessed using the RNA Nano 6000 Assay Kit of the Agilent Bioanalyzer 2100 system (Agilent Technologies, CA, USA). A total amount of 1 μg RNA per sample was used as input material for RNA sample preparations. Sequencing libraries were generated using NEBNext UltraTM RNA Library Prep Kit for Illumina (NEB, USA) following manufacturer’s recommendations and index codes were added to attribute sequences to each sample. Briefly, mRNA was purified from total RNA using poly-T oligo-attached magnetic beads. Fragmentation was carried out using divalent cations under elevated temperature in NEBNext First Strand Synthesis Reaction Buffer (5X). First strand cDNA was synthesized using random hexamer primer and M-MuLV Reverse Transcriptase. Second strand cDNA synthesis was subsequently performed using DNA Polymerase I and RNase H. Remaining overhangs were converted into blunt ends via exonuclease/polymerase activities. After adenylation of 3’ ends of DNA fragments, NEBNext Adaptor with hairpin loop structure were ligated to prepare for hybridization. To select cDNA fragments of preferentially 240 bp in length, the library fragments were purified with AMPure XP system (Beckman Coulter, USA). Then 3 μl USER Enzyme (NEB, USA) was used with size-selected, adaptor-ligated cDNA at 37°C for 15 min followed by 5 min at 95°C. PCR was then performed with Phusion High-Fidelity DNA polymerase, Universal PCR primers, and Index (X) Primer. At last, PCR products were purified (AMPure XP system) and library quality was assessed on the Agilent Bioanalyzer 2100 system.

#### Clustering, sequencing, and mapping

Clustering of index-coded samples was performed on a cBot Cluster Generation System using TruSeq PE Cluster Kit v4-cBot-HS (Illumina) according to the manufacturer’s instructions. After cluster generation, the library preparations were sequenced on an Illumina platform and paired-end reads were generated. The RNA sequencing was completed by Beijing Biomarker Co., Ltd. Raw reads of fastq format were processed through in-house perl scripts. At this step, clean reads were obtained by removing reads that contain adapter or ploy-N, and low-quality reads (N proportion > 10%, the number of bases with Q ≤ (N proportion > 10%, the number of bases wi) from raw data. In this parameter, Q represents the Quality Score which is integer mapping of the probability of base calling error. Q-value comes from Phred base quality formula: Q = −10 ∗ log_10_P in which P is the probability of base recognition error ^[56]^. At the same time, Q20, Q30, GC-content, and sequence duplication level of the clean data were calculated. All subsequent analyses were based on clean data with high quality. These clean reads were then mapped to a reference genome sequence (*VdLs.17*) ^[41]^ with Hisat2 tools. Only reads with a perfect match or only one mismatch from the reference genome were further analyzed and annotated.

### Bioinformatic analysis

#### Gene expression normalization, DEG recognition and gene annotation

Gene expression levels were quantified by fragments per kilobase of transcript per million fragments mapped. The formula is shown as follow: FPKM = cDNA Fragments / {Mapped Fragments(Millions) × Transcript Length(kb)}. Differential expression analysis was performed using the DESeq2 R package ^[57]^. DESeq2 provides statistical routines for determining differential expression in digital gene expression data using a model based on the negative binomial distribution. The resulting P values were adjusted using the Benjamin and Hochberg’s approach for controlling the false discovery rate. Genes with |log2(fold change) | > 1 an adjusted P-value < 0.05 found by DESeq2 were assigned as differentially expressed. Gene function was annotated using the following databases: NCBI non-redundant (Nr) protein sequences, NCBI non-redundant nucleotide (Nt) sequences, Protein family (Pfam), KEGG Ortholog (KO) database, Gene Ontology (GO), and Swiss-Prot, a manually annotated and reviewed protein sequence database. GO term and KEGG pathway enrichment of target gene set analysis was implemented by ClusterProfiler R package (v3.16.0) ^[58]^. Transcription factors (TFs) were predicted by plantTFDB (v4.0) ^[59]^ and animalTFDB (v3.0) ^[60]^. PROMO database (v8.3) ^[61, 62]^ was used to identify the putative transcription factor binding sites (TFBS).

#### Co-expression analysis

Weighted gene co-expression network analysis (WGCNA) is a widely used bioinformatic tool that incorporates hierarchical clustering to identify modules with different tree-cutting thresholds through summarizing each module by the module eigengene or an intramodular hub gene ^[49]^. Only genes were considered for analysis that had at least for one time point over all three replicates fpkm > 0.1 and coefficient of variation (CV) > 0.3 were considered. Next, the WGCNA R package (v1.69) ^[49]^ was used to identify modules of highly correlated genes across all samples with the gene fpkm matrix. Connection strengths among genes were estimated on the basis of correlation of expression data, and clusters of highly interconnected genes were identified as modules ^[63]^. Module detection was performed with default merge-CutHeight = 0.15 and the minimal module size was set to 30 genes. Upon detection, highly correlated modules were merged using a cut height of 0.25. The module eigengene adjacency heatmap was drawn using the data calculated with Pearson and Bicor (Fig. 4A, B). For each module, the expression profile of the module eigengene was calculated, which is defined as the first principal component of the module’s expression data. To examine the gene expression pattern of clustered modules, line graphs were drawn with growing phases on the X - axis and eigengene expression value on the Y - axis. GO and KEGG enrichments were performed for each selected module. For each gene, the intramodular connectivity (kME) was calculated, which represents Pearson correlation of individual genes with respective module eigengenes ^[64]^.

#### Data visualization

For the enrichment analysis of individual modules, only genes were considered that had an intramodular connectivity (kME) of greater than 0.5. Tableau (v2020.1) was used for drawing enrichment bubble plot. The protein-protein interaction (PPI) network was output from string database (v11.0) ^[65]^ and visualized by cytoscape (v3.7.0). TBtools (v0.6696) ^[66]^ was utilized to draw a gene heatmap.

#### Quantitative Real-Time PCR (qRT-PCR)

For gene expression analysis via Quantitative Real-Time PCR (qRT-PCR), the H and M samples grown on cellulose-covered PDA plates for 3 d, 4 d, 5 d and 6 d, 8 d, 10 d were used for RNA extraction with *TransZol* Up Plus RNA Kit. Then, 1 - 2 mg of isolated total RNA was reverse-transcribed with oligo (dT) primers using the First-Strand Super Script III cDNA Synthesis Kit (Life Technologies). The qRT-PCR was performed in an ABI QuantStudio 5 using *TransStart* Top Green qPCRSuperMix (AQ131). Cycling conditions were 10 min at 95°C, followed by 10 s at 94°C, then 40 cycles of 10 s at 56°C and 10 s at 72°C (S1 Table). The *V. dahliae* β-tubulin (*Vdβt*) ^[67]^ was used as a control. Primers were listed in (S2 Table). The arithmetic mean of reference gene Ct values served as normalization factor. For the fold change (FC) expression of qRT-PCR data, the gene expression in *Vd_BP_2_* strain served as control and the 2^-△△Ct^ method was used ^[68]^.

## Acknowledgements

The authors thank Biomarker Technology Co. (Beijing, China) for RNA-Seq and for providing analytical platform (BMKCloud). We thank Hangzhou Cambrian Biotechnology Co. Ltd. for performing real-time PCR of differentially express genes (DEGs).

## Supplementary information

**S1.a Fig.** Kyoto encyclopedia of genes and genomes (KEGG) enrichment of differentially expressed gene (DEG) sets with q(FDR) < 0.05.

**S1.b Fig.** KEGG enrichment of directionally expressed DEG sets with q(FDR) < 0.05.

**S2 Fig.** Sample clustering and soft power threshold. (a) Sample clustering with weighted gene correlation network analysis (WGCNA) to detect outliers. Td: defoliating strain *V. dahliae_086*; Tn: non-defoliating strain *V. dahliae_BP_2_*; H: Hyphal growth stage; M: Microsclerotia production stage; S: Spore germination stage. (b) Comparison of scale-free topology model and mean connectivity to pick soft power threshold. Red line was drawn at R^2^ = 0.8.

**S1 Table** qRT-PCR reaction mix.

**S2 Table** Primer sequences for qRT-PCR

**S3.a Table** GO enrichment of directionally expressed DEG sets with p < 0.005

**S3.b Table** KEGG enrichment of directionally expressed DEG sets with q (FDR) < 0.05

## References

1. Pegg G. The impact of Verticillium diseases in agriculture. Phytopathologia Mediterranea. 1984;23(2/3):176–192.

2. Puhalla J, Hummel M. Vegetative compatibility groups within *Verticillium dahliae*. Phytopathology. 1983;73(9):1305–1308.

3. El-Zik KM. Integrated control of Verticillium wilt of cotton. Plant Disease. 1985;69(12):1025–1032.

4. Hu XP, Gurung S, Short DP, Sandoya GV, Shang WJ, Hayes RJ, et al. Nondefoliating and defoliating strains from cotton correlate with races 1 and 2 of *Verticillium dahliae*. Plant disease. 2015;99(12):1713–1720.

5. Klebahn H. Beiträge zur Kenntnis der Fungi imperfecti: éditeur non identifié. 1914.

6. Duy P, Le AG, Thao T. Tran, Rodney J. Co-occurrence of defoliating and non-defoliating pathotypes of *Verticillium dahliae* in field-grown cotton plants in New South Wales, Australia. Plants. 2020;9:750.

7. Paplomatas E, Bassett D, Broome J, DeVay J. Incidence of Verticillium wilt and yield losses of cotton cultivars (*Gossypium hirsutum*) based on soil inoculum density of *Verticillium dahliae*. Phytopathology. 1992;82(12):1417–1420.

8. Perry JW, Evert RF. Structure of microsclerotia of *Verticillium dahliae* in roots of ‘Russet Burbank’ potatoes. Canadian journal of botany. 1984;62(2):396–401.

9. Zhang Q, Gao X, Ren Y, Ding X, Qiu J, Li N, et al. Improvement of Verticillium wilt resistance by applying arbuscular mycorrhizal fungi to a cotton variety with high symbiotic efficiency under field conditions. International journal of molecular sciences. 2018;19(1):241.

10. Zhang F, Li X, Zhu S, Ojaghian MR, Zhang J. Biocontrol potential of *Paenibacillus polymyxa* against *Verticillium dahliae* infecting cotton plants. Biological Control. 2018;127:70–77.

11. Yang C, Guo W, Li G, Gao F, Lin S, Zhang T. QTLs mapping for Verticillium wilt resistance at seedling and maturity stages in *Gossypium barbadense L*. Plant Science. 2008;174(3):290–298.

12. Pajerowska-Mukhtar KM, Emerine DK, Mukhtar MS. Tell me more: roles of NPRs in plant immunity. Trends in plant science. 2013;18(7):402–411.

13. Parkhi V, Kumar V, Campbell LAM, Bell AA, Rathore KS. Expression of arabidopsis *NPR1* in transgenic cotton confers resistance to non-defoliating isolates of *Verticillium dahliae* but not the defoliating isolates. Journal of Phytopathology. 2010;158(11-12):822–825.

14. Gao X, Wheeler T, Li Z, Kenerley CM, He P, Shan L. Silencing *GhNDR1* and *GhMKK2* compromises cotton resistance to Verticillium wilt. The Plant Journal. 2011;66(2):293–305.

15. Li TG, Wang BL, Yin CM, Zhang DD, Wang D, Song J, et al. The *Gossypium hirsutum* TIR-NBS-LRR gene *GhDSC1* mediates resistance against Verticillium wilt. Molecular plant pathology. 2019;20(6):857–876.

16. Gong Q, Yang Z, Wang X, Butt HI, Chen E, He S, et al. Salicylic acid-related cotton (*Gossypium arboreum*) ribosomal protein *GaRPL18* contributes to resistance to *Verticillium dahliae*. BMC plant biology. 2017;17(1):59.

17. Yang Y, Chen T, Ling X, Ma Z. Gbvdr6, a gene encoding a receptor-like protein of cotton (*Gossypium barbadense*), confers resistance to Verticillium wilt in Arabidopsis and upland cotton. Frontiers in plant science. 2018;8:2272.

18. Li C, He Q, Zhang F, Yu J, Li C, Zhao T, et al. Melatonin enhances cotton immunity to Verticillium wilt via manipulating lignin and gossypol biosynthesis. The Plant Journal. 2019;100(4):784–800.

19. Aro N, Pakula T, Penttilä M. Transcriptional regulation of plant cell wall degradation by filamentous fungi. FEMS microbiology reviews. 2005;29(4):719–739.

20. Houston K, Tucker MR, Chowdhury J, Shirley N, Little A. The plant cell wall: a complex and dynamic structure as revealed by the responses of genes under stress conditions. Frotiers in plant science. 2016;7(984).

21. Sangeeta Y, Yadav P, Dinesh Y, Yadav K. Pectin lyase: a review. Process Biochemistry. 2009;44(1):1–10.

22. Roeder DL, Collmer A. Marker-exchange mutagenesis of a pectate lyase isozyme gene in *Erwinia chrysanthemi*. Journal of bacteriology. 1985;164(1):51–56.

23. Yang Y, Zhang Y, Li B, Yang X, Dong Y, Qiu D. A *Verticillium dahliae* pectate lyase induces plant immune responses and contributes to virulence. Frontiers in plant science. 2018;9:1271.

24. Ruiz-Roldán M, Di Pietro A, Huertas-González M, Roncero M. Two xylanase genes of the vascular wilt pathogen *Fusarium oxysporum* are differentially expressed during infection of tomato plants. Molecular General Genetics. 1999;261(3):530–536.

25. Cooper R: The role of cell wall-degrading: Enzymes in infection and damage. Plant disease: Infection, damage loss. 1984:13–27.

26. Liu Q, Talbot M, Llewellyn D. Pectin methylesterase and pectin remodelling differ in the fibre walls of two gossypium species with very different fibre properties. PLoS One. 2013;8(6).

27. Xu JR, Hamer JE. MAP kinase and cAMP signaling regulate infection structure formation and pathogenic growth in the rice blast fungus *Magnaporthe grisea*. Genes development. 1996;10(21):2696–2706.

28. Amihud Y, Lev B. Does corporate ownership structure affect its strategy towards diversification? Strategic Management Journal. 1999;20(11):1063–1069.

29. Mayorga ME, Gold SE. A MAP kinase encoded by the *ubc3* gene of *Ustilago maydis* is required for filamentous growth and full virulence. Molecular microbiology. 1999;34(3):485–497.

30. Zheng L, Campbell M, Murphy J, Lam S, Xu JR. The *BMP1* gene is essential for pathogenicity in the gray mold fungus *Botrytis cinerea*. Molecular Plant-Microbe Interactions. 2000;13(7):724–732.

31. Wang Y, Tian L, Xiong D, Klosterman SJ, Xiao S, Tian C. The mitogen-activated protein kinase gene, *VdHog1*, regulates osmotic stress response, microsclerotia formation and virulence in *Verticillium dahliae*. Fungal Genetics Biology. 2016;88:13–23.

32. Tian L, Wang Y, Yu J, Xiong D, Zhao H, Tian C. The mitogen-activated protein kinase kinase *VdPbs2* of *Verticillium dahliae* regulates microsclerotia formation, stress response, and plant infection. Frontiers in microbiology. 2016;7:1532.

33. Fan R, Klosterman SJ, Wang C, Subbarao KV, Xu X, Shang W, et al. *Vayg1* is required for microsclerotium formation and melanin production in *Verticillium dahliae*. Fungal Genetics Biology. 2017;98:1–11.

34. Louw C, Young PR, Van Rensburg P, Divol B. Regulation of endo-polygalacturonase activity in Saccharomyces cerevisiae. 2009;10(1):44–57.

35. Kubicek CP, Starr TL, Glass NL. Plant cell wall-degrading enzymes and their secretion in plant-pathogenic fungi. Annual review of phytopathology. 2014;52:427–451.

36. Kombrink A, Rovenich H, Shi-Kunne X, Rojas-Padilla E, van den Berg GC, Domazakis E, et al. *Verticillium dahliae LysM* effectors differentially contribute to virulence on plant hosts. Molecular plant pathology. 2017;18(4):596–608.

37. De Jonge R, Thomma BP: Fungal LysM effectors: extinguishers of host immunity? Trends in microbiology. 2009;17(4):151–157.

38. Dürrenberger F, Wong K, Kronstad JW. Identification of a cAMP-dependent protein kinase catalytic subunit required for virulence and morphogenesis in *Ustilago maydis*. Proceedings of the National Academy of Sciences. 1998; 95(10):5684–5689.

39. Tzima A, Paplomatas EJ, Rauyaree P, Kang S. Roles of the catalytic subunit of cAMP-dependent protein kinase A in virulence and development of the soilborne plant pathogen *Verticillium dahliae*. Fungal Genetics Biology. 2010;47(5):406–415.

40. Bejarano-Alcázar J, Melero-Vara JM, Blanco-López MA, Jiménez-Díaz RM. Influence of inoculum density of defoliating and nondefoliating pathotypes of *Verticillium dahliae* on epidemics of Verticillium wilt of cotton in southern Spain. Phytopathology. 1995;85(12):1474–1481.

41. Klosterman SJ, Subbarao KV, Kang S. Comparative genomics yields insights into niche adaptation of plant vascular wilt pathogens. PLoS pathogen. 2011;7(7):e1002137.

42. Pan T, Coleman JE. GAL4 transcription factor is not a “zinc finger” but forms a Zn (II) 2Cys6 binuclear cluster. 1990;87(6):2077–2081.

43. Ekaterina S. Transcription factors in fungi: TFome dynamics, three major families, and dual-specificity TFs. Frontiers in genetics. 2017;8:53.

44. Ahmad S, Mohammed M, Mekala LP, Chintalapati S, Chintalapati VR. Tryptophan, a non-canonical melanin precursor: New L-tryptophan based melanin production by *Rubrivivax benzoatilyticus* JA2. Scientific Reports. 2020;10(1):1–12.

45. Carrasco M, Rozas JM, Alcaíno J, Cifuentes V, Baeza M. Pectinase secreted by psychrotolerant fungi: identification, molecular characterization and heterologous expression of a cold-active polygalacturonase from *Tetracladium sp*. Microbial cell factories. 2019;18(1):45.

46. Kirkman-Correia C, Stroke I, Fields S. Functional domains of the yeast *STE12* protein, a pheromone-responsive transcriptional activator. Molecular Cellular Biology. 1993;13(6):3765–3772.

47. Lo WS, Dranginis AM. The cell surface flocculin *Flo11* is required for pseudohyphae formation and invasion by *Saccharomyces cerevisiae*. Molecular biology of the cell. 1998;9(1):161–171.

48. Heale JB; Isaac I. Environmental factors in the production of dark resting structures in *Verticillium alboatrum*, *V. dahliae* and *V. tricorpus*. Transactions of the British Mycological Society. 1965;48(1):39–50.

49. Langfelder P, Horvath S. WGCNA. an R package for weighted correlation network analysis. BMC bioinformatics. 2008;9(1):559.

50. Tzima AK, Paplomatas EJ, Rauyaree P, Ospina-Giraldo MD, Kang S. *VdSNF1*, the sucrose nonfermenting protein kinase gene of *Verticillium dahliae*, is required for virulence and expression of genes involved in cell-wall degradation. Molecular Plant-Microbe Interactions. 2011;24(1):129–142.

51. Have A, Tenberge KB, Benen JA, Tudzynski P, Visser J, van Kan JA: The contribution of cell wall degrading enzymes to pathogenesis of fungal plant pathogens. In: Agricultural Applications. Springer; 2002:341–358.

52. Gibson DM, King BC, Hayes ML, Bergstrom GC. Plant pathogens as a source of diverse enzymes for lignocellulose digestion. Current opinion in microbiology. 2011;14(3):264–270.

53. Sun X, Lu X, Zhang J, Zhu S. Identification of phytotype of *Verticillium dahliae* isolates on cotton in Zhejiang province and phenotypic analysis on inhibitory effect by high tempreture. Journal of Zhejiang University. 2016;42(6):671–678.

54. Gao F, Zhou BJ, Li GY, Jia PS, Li H, Zhao YL, et al. A glutamic acid-rich protein identified in *Verticillium dahliae* from an insertional mutagenesis affects microsclerotial formation and pathogenicity. PloS one. 2010;5(12).

55. Cassago A, Panepucci RA, Baião AMT, Henrique-Silva FJBm. Cellophane based mini-prep method for DNA extraction from the filamentous fungus Trichoderma reesei. 2002;2(1):14.

56. Ewing B, Hillier L, Wendl MC, Green P. Base-calling of automated sequencer traces usingPhred. I. Accuracy assessment. Genome research. 1998;8(3):175–185.

57. Love MI, Huber W, Anders S. Moderated estimation of fold change and dispersion for RNA-seq data with DESeq2. Genome biology. 2014;15(12):550.

58. Yu G, Wang LG, Han Y, He QY. clusterProfiler: an R package for comparing biological themes among gene clusters. Omics: a journal of integrative biology. 2012;16(5):284–287.

59. Jin J, Tian F, Yang DC, Meng YQ, Kong L, Luo J, et al: PlantTFDB 4.0: toward a central hub for transcription factors and regulatory interactions in plants. Nucleic acids research. 2016:gkw982.

60. Hu H, Miao Y, Jia L, Yu Q, Zhang Q, Guo A. AnimalTFDB 3.0: a comprehensive resource for annotation and prediction of animal transcription factors. Nucleic acids research. 2019;47(D1):D33–D38.

61. Messeguer X, Escudero R, Farré D, Nuñez O, Martínez J, Albà MM. PROMO: detection of known transcription regulatory elements using species-tailored searches. Bioinformatics. 2002;18(2):333–334.

62. Farré D, Roset R, Huerta M, Adsuara JE, Roselló L, Albà MM, Messeguer X. Identification of patterns in biological sequences at the ALGGEN server: PROMO and MALGEN. Nucleic acids research. 2003;31(13):3651–3653.

63. Hu GJ, Hovav R, Grover CE, Faigenboim-Doron A, Kadmon N, Page JT, et al. Evolutionary conservation and divergence of gene coexpression networks in Gossypium (cotton) seeds. Genome Biology and Evolution. 2017;8(12):3765–3783.

64. Lanver D, Müller AN, Happel P, Schweizer G, Haas FB, Franitza M, et al. The biotrophic development of *Ustilago maydis* studied by RNA-seq analysis. The Plant Cell. 2018;30(2):300–323.

65. Szklarczyk D, Gable AL, Lyon D, Junge A, Wyder S, Huerta-Cepas J, et al. STRING v11: protein-protein association networks with increased coverage, supporting functional discovery in genome-wide experimental datasets. Nucleic Acids Res. 2019;47:D607–613. https://string-db.org/

66. Chen C, Chen H, He Y, Xia R. TBtools, a toolkit for biologists integrating various biological data handling tools with a user-friendly interface. BioRxiv. 2018:289660.

67. Atallah ZK, Bae J, Jansky SH, Rouse DI, Stevenson WR. Multiplex real-time quantitative PCR to detect and quantify *Verticillium dahliae* colonization in potato lines that differ in response to Verticillium wilt. Phytopathology. 2007;97(7):865–872.

68. Livak KJ, Schmittgen TD. Analysis of relative gene expression data using real-time quantitative PCR and the 2^-ΔΔCT^ method. Methods. 2001;25(4):402–408.

